# Estrogen deprivation triggers an immunosuppressive phenotype in breast cancer cells

**DOI:** 10.1101/715136

**Authors:** Daniela Hühn, Pablo Martí-Rodrigo, Silvana Mouron, Catherine S. Hansel, Kirsten Tschapalda, Bartlomiej Porebski, Maria Häggblad, Louise Lidemalm, Miguel A. Quintela-Fandino, Jordi Carreras-Puigvert, Oscar Fernandez-Capetillo

**Affiliations:** Science for Life Laboratory, Division of Genome Biology, Department of Medical Biochemistry and Biophysics, Karolinska Institute, S-171 21 Stockholm, Sweden; Breast Cancer Clinical Research Unit, Spanish National Cancer Research Centre (CNIO), Madrid 28029, Spain; Genomic Instability Group, Spanish National Cancer Research Centre (CNIO), Madrid 28029, Spain

## Abstract

Estrogen receptor (ER)-positive breast tumors are routinely treated with estrogen-depriving therapies. Despite their effectiveness, patients often progress into a more aggressive form of the disease. Through a chemical screen oriented to identify chemicals capable of inducing the expression of the immune-checkpoint ligand PD-L1, we found antiestrogens as hits. Subsequent validations confirmed that estrogen deprivation or ERα depletion induces PD-L1 expression in ER-positive breast cancer cells, both *in vitro* and *in vivo*. Likewise, PD-L1 expression is increased in metastasis arising from breast cancer patients receiving adjuvant hormonal therapy for their local disease. Transcriptome analyses indicate that estrogen deprivation triggers a broad immunosuppressive program, not restricted to PD-L1. Accordingly, estrogen deprived MCF7 cells are resistant to T-cell mediated cell killing, in a manner that can be reverted by estradiol. Our study reveals that while antiestrogen therapies effectively limit tumor growth in ER-positive breast cancers, they also trigger a transcriptional program that favors immune evasion.

## INTRODUCTION

Breast cancer (BC) is the most frequent cancer in women worldwide and the second-cause of cancer-related mortality (Bray, Ferlay et al., 2018). Around 75% of BC cases are tumors that express Estrogen Receptor alpha (ERα) and are dependent on its transcriptional activity for survival. Accordingly, therapies directed to limit ERα signaling such as ER antagonists (e.g. tamoxifen) or ER downregulators (e.g. fulvestrant) constitute the first-line treatment for ER^+^ BC (reviewed in (Siersbaek, Kumar et al., 2018)). Specifically, the current standard of care is a combination of the hormone therapy with cyclin-dependent kinase 4/6 (CDK4/6) inhibitors, even though, until very recently, clinical trials had failed to see a significant increase in overall survival with this combination (Im, Lu et al., 2019). In any case, and while antiestrogen therapy is certainly effective, around one fifth of the patients relapse into a metastatic stage for which there is no cure. Because of this, there is an intensive effort in testing the efficacy of new therapies, to be used alone or in combination with hormone therapy, for the treatment of ER^+^ BC. Not surprisingly, there is particular interest in exploring the potential of therapies targeting immune checkpoints such as those mediated by cytotoxic-T-lymphocyte-associated antigen 4 (CTLA-4) and programmed death-1 (PD-1) (Wei, Duffy et al., 2018), although initial evidence indicate that ER^+^ tumors are not very responsive due to low numbers of infiltrating lymphocytes and a low mutational burden (Esteva, Hubbard-Lucey et al., 2019).

Estrogens, most frequently 17β-estradiol (E2), have widespread effects on transcription which are to a large extent mediated by binding to two members of the nuclear receptor family, ERα (*ESR1*) and ERβ (*ESR2*). Upon binding to estrogens, these factors homodimerize and bind to target sequences on chromatin where they regulate transcription preferentially at distal enhancers (Hah, Murakami et al., 2013, Li, Notani et al., 2013). Besides their well-known roles in reproductive organs, estrogens have generalized effects in other tissues including bone, liver, colon, adipose tissue, kidney, skin and the cardiovascular and central nervous systems (Eyster, 2016). In addition, emerging evidence indicates a particularly important role of estrogens in suppressing inflammation, which helps to explain the gender-related differences found in diseases such as multiple sclerosis or rheumatoid arthritis that are alleviated during pregnancy when estrogen levels peak (Confavreux, Hutchinson et al., 1998, Nelson & Ostensen, 1997). On the contrary, the use of oral contraceptives reducing sex hormone levels has been shown to aggravate inflammatory bowel disease (Khalili, Higuchi et al., 2013). This immunosuppressive role of estrogens also plays an important role in cancers associated to inflammation such as hepatocellular carcinoma, which is three to five times more frequent in men than in women (Naugler, Sakurai et al., 2007). However, the contribution of this phenomenon to BC onset and response to hormone-therapy is still poorly understood. Here, by screening for molecules that modulate PD-L1 expression, we reveal a role for estrogens in suppressing a broad inflammatory transcriptional program in ER^+^ BC cells, which limits their clearance by the immune system.

## RESULTS

### A chemical screen to identify regulators of PD-L1 expression

In order to identify chemicals capable of regulating surface PD-L1 expression, we conducted a High-Throughput screen using 4,216 pharmacologically active compounds (**Fig. 1A**; see Methods for details). The screening was conducted in the human lung cancer cell line A549, which was previously shown to express PD-L1 upon interferon gamma (IFNγ) stimulation (Stanciu, Bellettato et al., 2006). Since we were originally more focused on discovering downregulators, the screening was conducted on A549 cells that were previously treated with 100 ng/ml of IFNγ for 24 hrs and then subsequently with the compounds from the library for another 24 hrs. At this point, cells were stained with anti-PD-L1 antibodies, fixed and processed for High-Throughput Microscopy (HTM). As expected, wells treated with only IFNγ (positive control) showed a significant increase of PD-L1 expression when compared with DMSO-treated negative controls. In this experimental setup, corticosteroids were the most enriched compound class found among the downregulators (**Fig. 1B** and **Table S1**). Subsequent validation experiments confirmed that 3 independent corticosteroids (dexamethasone, hydrocortisone and prednisone) significantly reduced the surface levels of PD-L1 in IFNγ-treated A549 cells (**Fig. S1**). Besides corticosteroids, inhibitors of the Janus kinases (JAK1/2) were also found among the downregulators, which are known to be involved in the IFNγ-dependent induction of PD-L1 (Hao, Chapuy et al., 2014). Noteworthy, this work can help to understand recent reports that either *JAK1/2* mutations (Shin, Zaretsky et al., 2017) or baseline corticosteroid treatment (Arbour, Mezquita et al., 2018, Gourd, 2018) confer resistance to anti-PD-L1 therapy.

**Figure 1.**
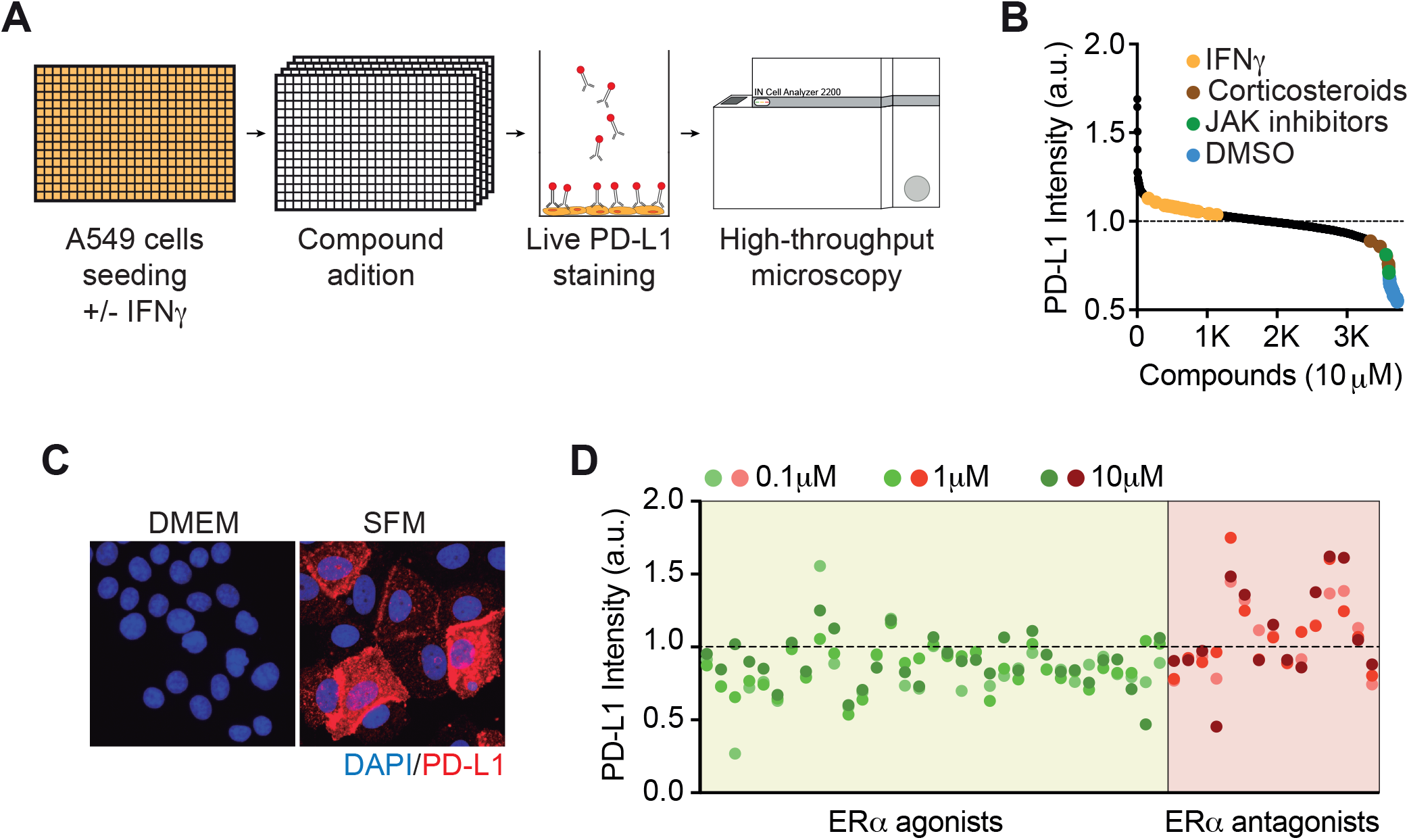
Small molecule screen for modulators of PD-L1 expression. **A,** Overview of the phenotypic screen workflow. Briefly, A549 cells were seeded in 100ng/ml IFNγ 24hrs before addition of 4216 compounds at 10μM. After 24hrs of compound exposure, cells were stained with an anti-PD-L1 antibody conjugated to phycoerythrin and fixed with formaldehyde. Nuclei were stained with Hoechst and the immunofluorescences were analysed by high-throughput microscopy (HTM). **B,** Hit distribution of the screen described in **A** illustrating the enrichment of JAK inhibitors and corticosteroids among the compounds reducing PD-L1 signal in HTM. PD-L1 levels in wells only stimulating with IFNγor negative controls (DMSO) are also shown. **C,** Immunofluorescence of PD-L1 (red) in MCF7 cells grown in normal or steroid-free medium (SFM) for 15 days. DAPI (blue) was used to stain DNA. **D,** Scatterplot of PD-L1 intensity levels in SFM-grown MCF7 cells treated with ERαagonists or antagonists, screened at three concentrations, 0.1, 1.0 and 10*μ*M.

With regard to up-regulators, there were very few molecules capable of inducing PD-L1 beyond the levels observed with IFNγ, which we failed to confirm in subsequent validation experiments (in many cases the hits were related to autofluorescence of the compounds). Nevertheless, we were intrigued by the presence of the selective estrogen receptor degrader (SERD) fulvestrant among this list of hits, even if the screening was done in A549 cells, given that previous reports have indicated reduced lung cancer mortality risk in BC patients treated with antiestrogens (Bouchardy, Benhamou et al., 2011), and that fulvestrant has actually been shown to reduce the growth of A549 xenografts (Marquez-Garban, Chen et al., 2007). In order to further investigate the impact of anti-estrogen therapies on PD-L1 expression we switched to MCF7, which is a widely used ER+ BC cell line. First, in order to evaluate the effect of a complete hormone deprivation, we grew MCF7 cells in steroid-free media (SFM) for two weeks. This led to a clear upregulation of PD-L1, which as expected was present on the cell membrane (**Fig. 1C**). Using this experimental setup, we conducted a secondary chemical screen, where we tested the effects of 25 ER agonists and 11 ER antagonists in a dose-response (**Fig. 1D**, **Table S2**). Overall, ER agonists reduced and antagonists further increased PD-L1 expression in SFM-grown MCF7 cells.

### ERα signaling suppresses PD-L1 expression in ER^+^ BC cells

To further evaluate the effect of estrogen signaling in ER^+^ BC cells, MCF7 cells were either hormone-deprived in SFM or treated with fulvestrant for 2 weeks. While both conditions led to a clear upregulation of surface PD-L1 levels as detected by flow cytometry, only the effect of the SFM-treatment could be reverted with ethinylestradiol (EE) (**Fig. 2A-D**). Equivalent results were observed by microscopy (**Fig. 2E**). The lack of effect of EE in downregulating PD-L1 expression in fulvestrant-treated MCF7 cells suggests that the observed effects are due to ERα, as the compound leads to the degradation of this receptor. Consistently, western blot analyses confirmed that EE reverted the increase in PD-L1 expression in SFM- but not fulvestrant-treated MCF7 cells, as the later lacked ERα expression (**Fig. 2F**). In addition, while SFM- or fulvestrant treatment induced PD-L1 expression in ERα^+^ cells, it failed to do so in ERα^−^ ones (**Fig. S2**). Quantitative reverse transcription polymerase chain reaction (qRT-PCR) analyses revealed that the upregulation of PD-L1 (*CD274*) levels in SFM- or fulvestrant-treated MCF7 cells occurred at the level of transcription (**Fig. 2G,H**). Finally, RNA interference-mediated downregulation of ERα (*ESR1*) but not ERβ (*ESR2*) also led to the upregulation of surface PD-L1 levels in MCF7 cells (**Fig. S3**). Collectively, these experiments reveal that estrogens suppress PD-L1 expression through the stimulation of ERα signaling.

**Figure 2.**
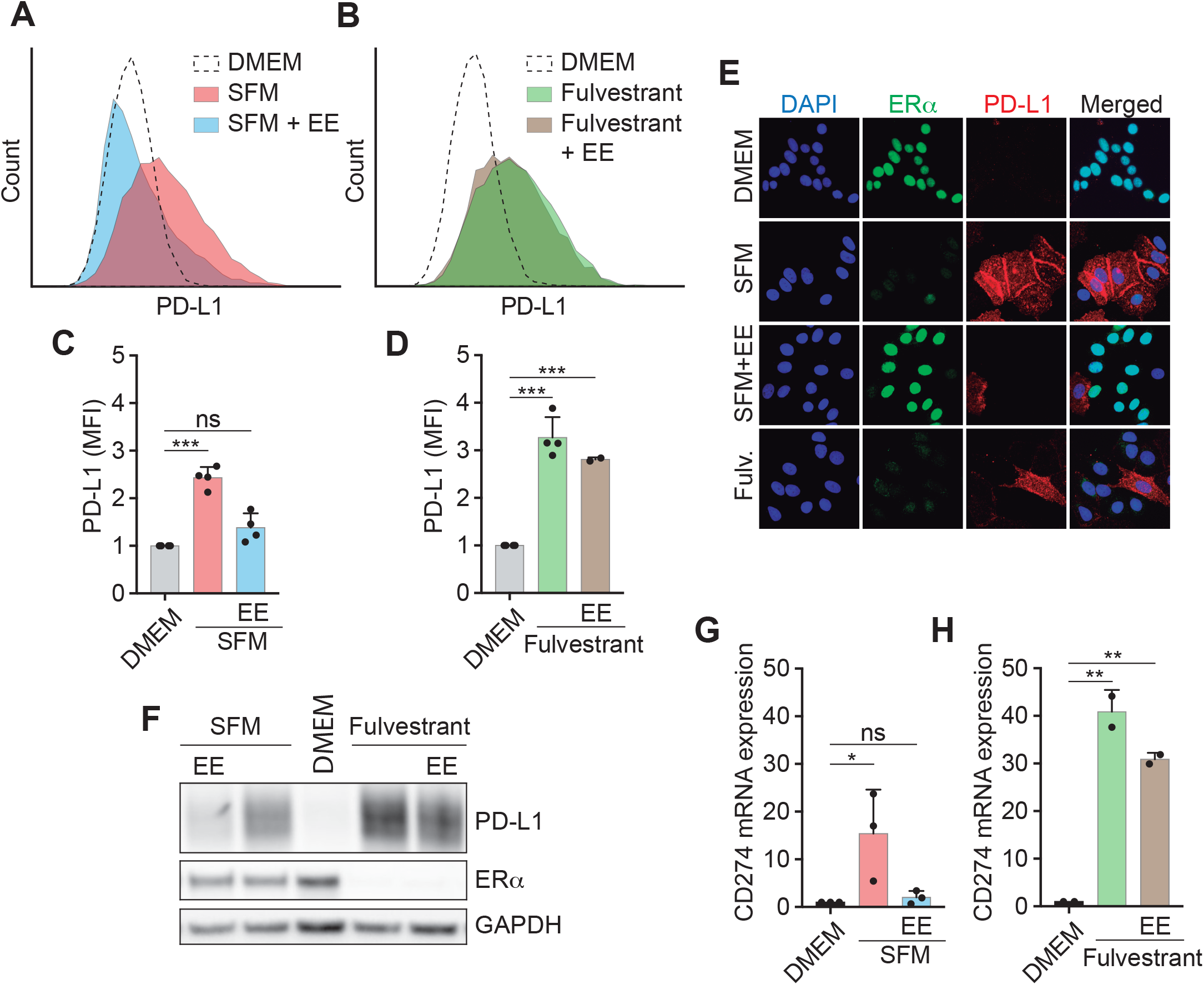
Estrogen-dependent suppression of PD-L1 expression in ER+ BC cells. **A,** Flow cytometry mediated assessment of surface PD-L1 levels in MCF7 cells grown DMEM, SFM or SFM plus EE for 14 days. EE (10 nM) was added for the last 3 days. **B,** Flow cytometry mediated assessment of surface PD-L1 levels in MCF7 cells grown DMEM or fulvestrant (1μM) for 14 days. EE (10 nM) was added for the last 3 days. **C,D** Quantification from the data shown in **A** (**C**) and **B** (**D**). **E,** Immunofluorescence illustrating ERα (green) and PD-L1 (red) levels in MCF-7 cells cultured in DMEM, SFM or fulvestrant (1 μM) for 20 days. Where indicated, EE (10 nM) was supplemented during the last 4 days. Representative images for each condition are shown. **F,** Western blot illustrating the levels of PD-L1 and ERα in MCF-7 cells grown in DMEM, SFM or DMEM containing 1μM fulvestrant (Fulv) for 14 days. Where indicated EE (10nM) was added for the final 3 days. GAPDH levels are shown for loading control. **G, H** qRT-PCR analysis of PD-L1 (*CD274*) expression in MCF-7 cells cultured in DMEM and SFM (**G**) or fulvestrant (1 μM, **H**) for 18 days. Where indicated, media was supplemented with EE (10 nM) for the last 3 days. *18S* RNA served as an internal control. \**p*<0,05; \*\**p*<0,01

### Estrogen deprivation triggers an inflammatory transcriptional program in MCF7 cells

To determine the mechanism by which estrogen suppresses PD-L1 expression in ER^+^ BC cells we analyzed how estrogen deprivation impacts the activation of Janus Kinase-Signal Transducer and Activator of Transcription proteins (JAK-STAT) and NF-κB signaling pathways, both of which are known regulators of PD-L1 expression and play important roles in the sensitivity to cancer immunotherapies (Manguso, Pope et al., 2017, Pan, Kobayashi et al., 2018, Peng, Hamanishi et al., 2015, Shin et al., 2017). In fact, a time-course of MCF7 cells grown in SFM revealed that both pathways were activated, as evidenced by the phosphorylation of STAT1 at Tyr 701 (p-STAT1^Tyr701^) and RelA (p65) at Ser 536 (p-p65^Ser536^), concomitantly to the upregulation of PD-L1 (**Fig. 3A**). Interestingly, addition of EE after 16 days in SFM for the last 4 days reverted STAT1 but not p65 phosphorylation, arguing that the JAK/STAT pathway is the primary mediator of the observed effects. Nevertheless, both the JAK2 inhibitor (CEP-33779) or the NF-κB inhibitor (CAPE) reduced the upregulation of PD-L1 induced by SFM in MCF7 cells (**Fig. 3B**).

**Figure 3.**
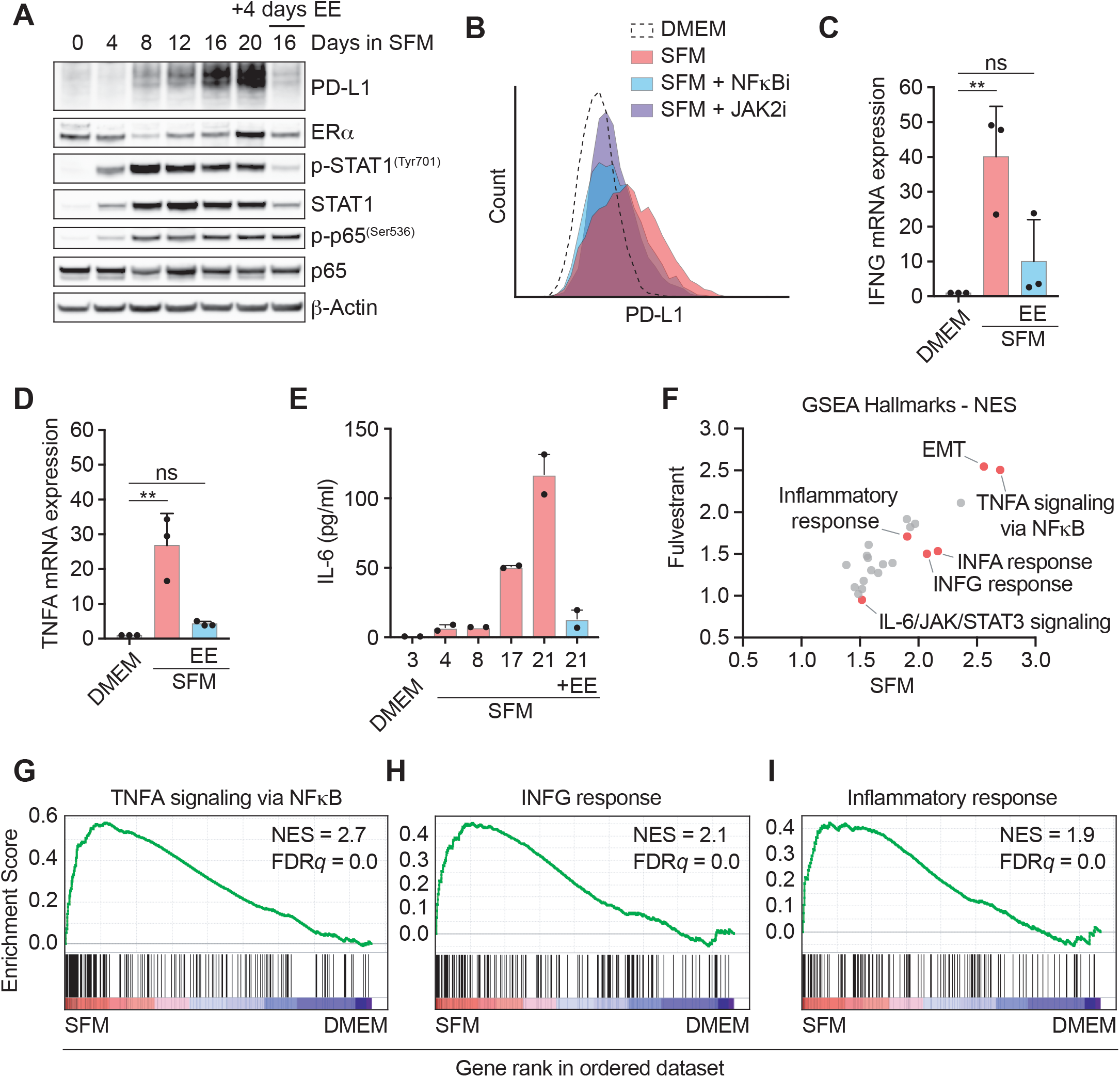
Estrogen signaling suppresses an inflammatory phenotype in MCF7 cells. **A,** Whole-cell lysates from MCF-7 cells cultured in SFM for the specified days were analyzed by western blotting using the indicated antibodies. Where indicated, 10nM EE was added at 16 days for the final 4 days. Total p65 and β-ACTIN served as loading controls. **B,** Flow cytometry mediated evaluation of PD-L1 membrane levels in MCF-7 cells grown in DMEM or SFM for 21 days, alone or in combination with the NFkB inhibitor CAPE (10μM) or JAK2 inhibitor CEP-33779 (10μ) for the last 3 days. **C,D** qRT-PCR analysis of IFNγ (*IFNG*) (**C**) or TNFα (*TNFA*) (**D**) mRNA levels in MCF-7 cells cultured as in (**A**). *18S* rRNA was used as an internal control. \*\**p*<0,01 **E,** Levels of IL-6 in the supernatant of MCF7 cells cultured in DMEM or SFM for the specified days as analyzed by LegendPlex-FACS (see Methods). Where indicated EE (10 nM) was added at day 17 for the last 4 days. **F,** GSEA Hallmark gene sets ranked by normalised enrichment score (NES) comparing the transcriptional programs triggered by 3 week treatments with fulvestrant (1 μM) or SFM in MCF7 cells, both normalised to DMEM. Selected hallmarks are indicated in red. **G,H,I** Pre-ranked GSEA on the genes from the hallmarks “TNFA signaling via NF-κB” (**G**), “IFNG response” (**H**) and “Inflammatory response” (**I**) obtained from RNAseq analysis comparing the transcriptome of MCF7 cells grown in DMEM or SFM for 3 weeks. The heatmap representation illustrates the overall upregulation of these pathways in estrogen-deprived MCF7 cells.

As to how JAK/STAT and NF-κB signaling are activated upon estrogen deprivation, we found increased levels of IFNγ and Tumor Necrosis Factor alpha (TNFα) in SFM-grown MCF7 cells, which are the primary cytokines involved in the activation of each pathway, respectively (**Fig. 3C, D**). Interestingly, and besides IFNγ, estrogen deprivation also induced the secretion of IL-6, which is a central inflammatory cytokine that stimulates JAK/STAT signaling and that is known to decrease the effectiveness of cancer immunotherapy (**Fig. 3E**) (Johnson, O’Keefe et al., 2018). Of note, given that estrogen deprivation arrests the growth of MCF7 cells and that IL-6 is also an important component of the senescence associated secretory phenotype (SASP) (Faget, Ren et al., 2019), we wondered whether PD-L1 expression was part of a SASP response in these cells. However, even if MCF7 cells grown in SFM showed several features of senescence such as increased levels of p21^Cip1^ and histone H3 lysine 9 trimethylation (H3K9me3) or an increased activity of the senescence-associated beta galactosidase (SA-βgal) (**Fig. S4A,B**), MCF7 cells induced to undergo senescence upon treatment with the p53 activator Nutlin-3 failed to upregulate PD-L1 (**Fig. S4C**).

To obtain a general view of the transcriptional changes induced by estrogen deprivation in ER^+^ BC cells, we conducted RNA sequencing (RNAseq) in MCF7 cells grown in SFM or with fulvestrant for 3 weeks. Analysis of Gene Set Enrichment Analysis (GSEA) hallmarks showed a good correlation between the transcriptional changes induced by both conditions (**Fig. 3F**). One of the common hallmarks was the Epithelial-Mesenchymal Transition (EMT), which is consistent with the change in morphology that is observed with these treatments. Moreover, and in support to the previous data, “TNFα signaling via NF-κB”, “IFNγ response” or “Inflammatory response” were amongst the most significantly induced hallmarks (**Fig, 3F-I** **and Fig. S5**). Thus, estrogen deprivation triggers a broad inflammatory transcriptional program in MCF7 cells, which includes the activation of known activators of PD-L1 expression such as NF-κB and IFNγ.

### Estrogen deprivation limits T-cell mediated cell killing of MCF7 cells

To evaluate the extent to which the transcriptional program triggered by estrogen deprivation in BC cells affected their sensitivity to immune cells, we conducted a T-cell mediated cell killing assay in MCF7 cells. To do so, MCF7 cells stably expressing mCherry fused to a nuclear localization sequence were co-cultured in the presence of activated primary T-cells and followed by live cell imaging for 3 days. Remarkably, MCF7 cells that were previously grown on SFM or with fulvestrant were significantly resistant to their killing by T-cells (**Fig. S6A**). In addition, EE was able to alleviate the effects of the SFM treatment and increased the elimination of MCF7 cells by T-cells. Equivalent results were obtained by measuring apoptosis through the use of a fluorescent caspase-3/7 target (**Fig. S6B**). As expected, no cell killing was observed when MCF7 cells were co-cultured in the presence of T-cells that were not previously activated (**Fig. S6C,D**). Thus, estrogen deprivation increases the resistance of ER^+^ MCF7 cells to their killing by T lymphocytes.

### ERα inversely correlates with PD-L1 expression in breast cancer

Recent analyses of The Cancer Genome Atlas (TCGA) project have indicated higher levels of PD-L1 in Triple Negative Breast Cancer (TNBC) when compared to non-TNBC subtypes (Mittendorf, Philips et al., 2014). Based on our findings, we explored whether this correlation could be linked to the expression of ERα. Indeed, gene expression analysis of 1904 BCs (ERα^+^: 1459; ERα^−^:445) from the METABRIC cohort of the TCGA dataset revealed significantly higher levels of PD-L1 mRNA expression (*CD274*) in the ERα^−^ cohort (**Fig. 4A**). Similar results could be observed at the protein level when comparing PD-L1 expression between 5 ERα^+^ cell lines (MCF-7, T47D, CAMA-1, ZR-75-1 and BT-474) and 3 ERα^−^ ones (MDA-MB-231, HCC1937 and BT-549) (**Fig. 4B**). Next, to determine if PD-L1 expression was also induced *in vivo* in response to an anti-estrogen treatment we used a transgenic mouse model of ER^+^ BC based on the expression of the Polyoma Virus middle T Antigen downstream of the mouse mammary tumor virus long terminal repeat (MMTV-PyMT) (Guy, Cardiff et al., 1992). In agreement with our *in vitro* findings, treatment of MMTV-PyMT transgenic mice harboring BC with the ER agonist tamoxifen led to a significant increase in PD-L1 expression in the tumors (**Fig. 4C,D**). Finally, we evaluated PD-L1 expression in tissue from a small cohort of human patients of ER^+^ BC (clinical and demographic characteristics detailed on **Table S3**) for which we obtained paired biopsies from the primary tumor and the metastases that emerged during or after adjuvant hormonal therapy. While PD-L1 expression was virtually absent in the primary tumors, areas of PD-L1 expressing cells could be detected in half of the metastatic samples **(Fig. 4E,F**).

**Figure 4.**
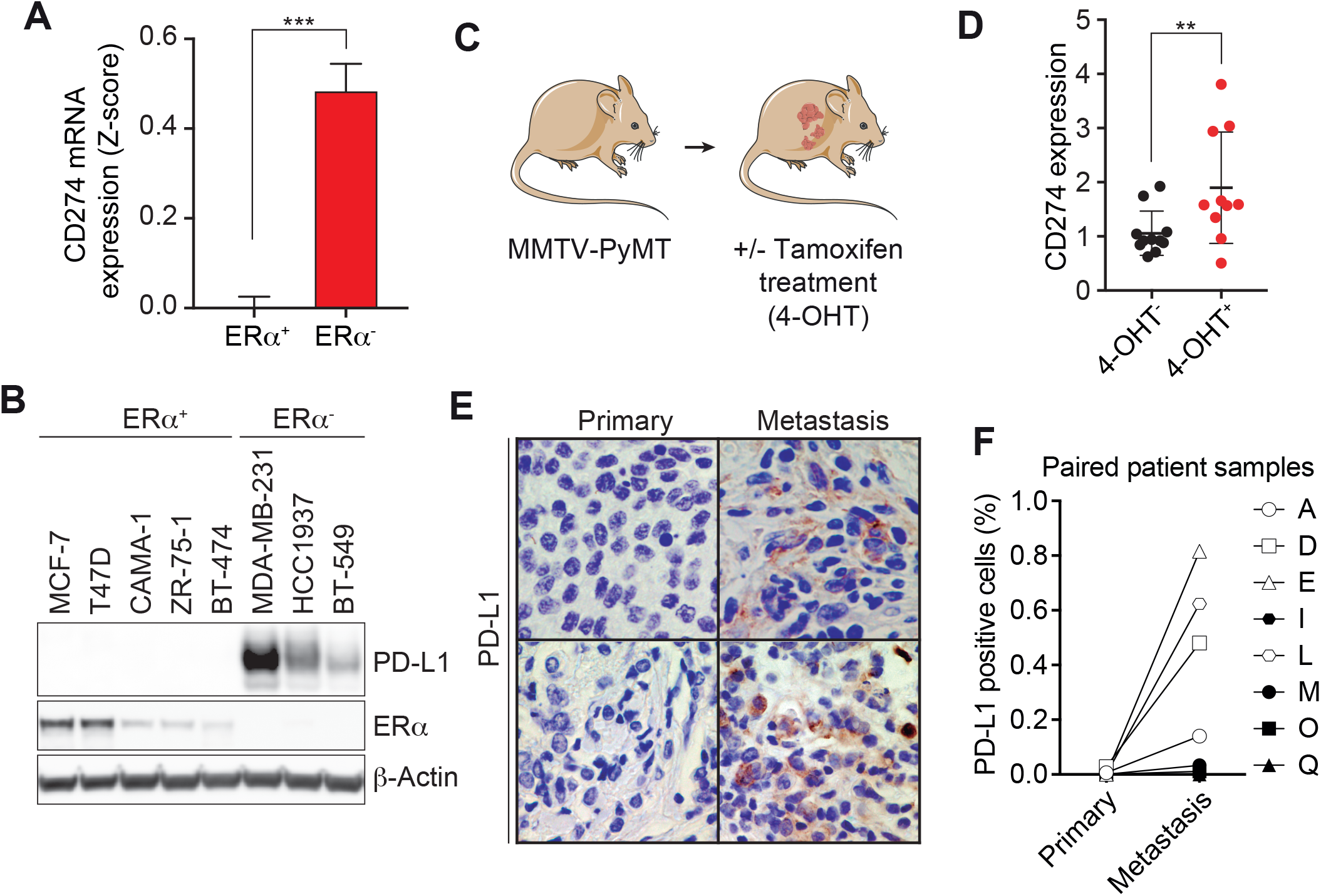
Inverse correlation between ERα and PD-L1 in BC. **A,** *CD274* mRNA expression levels in ERα^+^ (n=1459) and ERα^−^ (n=445) patient samples. Normalised Z-scores were extracted from the METABRIC breast cancer TCGA dataset (Pereira et al., 2016). A factor was added to both populations in order to set the ERα+ group to 0. Data are presented as mean +SEM and two-tailed Student’s t-test was used to calculate the statistical significance, ***p<0.001. **B,** Western blot analysis of PD-L1 and ERαprotein in a panel of ERα^+^ and ERα^−^ BC cell lines. β-Actin served as a loading control. **C,** Schematics of the experimental procedure in mice. MMTV-PyVT mice that spontaneously develop mammary tumors were treated daily from week 5 until with 4-hydroxytamoxifen (4-OHT) or vehicle. **D,** *Cd274* expression in tumors isolated from MMTV-PyVT mice treated with 4-OHT or vehicle as determined by RT-qPCR. Data were normalized to β-*Actin* mRNA levels. Graph represents the mean ±SD and two-tailed Student’s t-test was used to determine statistical significance, **p<0.01. **E,** Immunohistochemical staining of PD-L1 (brown) in paired samples of primary and metastatic human ER+ BC from two different patients. Sections were stained with Hematoxylin-Eosin to aid in the pathological analysis. **F**, Quantification of the percentage of PD-L1 positive cells in paired primary tumor and metastasis samples from 8 patients of ERα-positive BC.

## DISCUSSION

Estrogen was one of the first hormones to be described and for a while was thought to only act in the female reproductive system (Eyster, 2016). Later studies revealed that estrogen receptors were widely expressed in many organs, and that estrogens played pleotropic physiological roles beyond reproduction. Among these, several studies have indicated that estrogen levels influence the severity of diseases linked to inflammation, including cancer (Khalili et al., 2013, Naugler et al., 2007, Nelson & Ostensen, 1997). However, somewhat surprisingly, the relevance of the link between estrogen signaling and inflammation has not been particularly focused on ER^+^ BC, which is the cancer type where estrogen signaling plays a central role. As to how ER modulates inflammation, prior evidence had indicated a suppressive role of ERα in NFκB (Kalaitzidis & Gilmore, 2005). Consistent with our findings, a recent study also indicated a role for ERα in regulating PD-L1 transcription, although the mechanism was unknown (Liu, Shen et al., 2018). Here, we show that the impact of estrogen deprivation is not restricted to a specific effect of PD-L1 but rather triggers a broad inflammatory transcriptional program in ER^+^ BC cells, which includes the secretion of cytokines that activate NFκB signaling such as TNFα, but also additional cytokines such as IFNγ and IL-6 that trigger the activation of the JAK/STAT pathway. The immunosuppressive phenotype of estrogen-deprived ER^+^ BC cells includes the expression of immune checkpoints such as PD-L1, which limit their clearance by the immune system. This phenotype switch that occurs upon estrogen deprivation should be considered when considering cancer immunotherapy regimens for the treatment of ER^+^ BC. Of note, and besides estrogen signaling, our chemical screen also identified corticosteroids as the most prominent class of drugs that reduces IFNγ-induced PD-L1 expression, providing an explanation for recent observations that indicate a reduced efficiency of anti-PD1/PD-L1 immunotherapies in patients undergoing a baseline treatment with corticosteroids (Gourd, 2018).

BC has classically been considered as poorly responsive to immunotherapy due to initial failures in vaccination or cytokine treatments in the 1980’s and 90’s (Buzdar, Blumenschein et al., 1984, Miles, Roche et al., 2011), and due to the fact that ER^+^ BC has a low mutational burden and is therefore immunologically “cold”. Nevertheless, metastatic BC is still the second cause of cancer-related mortality worldwide and new therapies are urgently needed. In this context, and due to the impressive effects that have been seen in patients of cancers with dismal prognosis such as melanoma or lung cancer, the efficacy of cancer immunotherapy is now a very active area of clinical investigation in the treatment of BC. As of September 2018, there were 285 clinical trials related to immunotherapy in BC, 208 of which were related to targeting the PD-1/PD-L1 checkpoint (reviewed in (Esteva et al., 2019)). However, since hormone therapy is a safe and effective approach for the initial treatment of ER^+^ BC, clinical trials are mostly oriented to either TNBC or to metastatic disease of any subtype. Still, around one fifth of the ER^+^ BC patients undergoing hormone therapy progress into metastasis. Based on our work, we suggest that a combination of hormone therapy with therapies targeting PD-1 or PD-L1 for the treatment of primary ER^+^ BC might facilitate the clearance of the residual BC cells that persist even at low growth rates, and thus reduce the percentage of patients that progress into metastatic disease.

## METHODS

### Cell culture, transfection and chemicals

A549, MCF7, T47D, HCC1937 and MDA-MB-231 cell line identity was confirmed using short-tandem repeat profiling analysis by ATCC. Except HCC1937, which were cultured in RPMI-1640, all cell lines were cultured in Dulbecco’s modified Eagle’s medium supplemented with 10% fetal calf serum and penicillin/streptomycin (100 U/ml) at 37°C in a 5% CO_2_ humidified incubator. In experiments requiring hormone depletion, cells were cultured in phenol red-free DMEM supplemented with 2mM L-glutamine and 10% charcoal-dextran stripped fetal calf serum (Sigma-Aldrich, F6765), hereafter termed steroid-free media (SFM). Where indicated 10nM 17α-Ethynylestradiol (Sigma-Aldrich, E4876) was added to SFM. For fulvestrant treatments, cells were cultured in standard DMEM supplemented with 1μM fulvestrant (ICI 182,780, Tocris, 1047) for the indicated number of days. During all treatments the respective media was exchanged every 4 days. For siRNA transfections, cells were seeded in 6-well plates and transfected the next day with 30pmol of *ESR1* (Sigma-Aldrich, SASI_Hs01_00078592), *ESR2* (Dharmacon, L-003402-00-0005) or Ctrl (Dharmacon, D-001810-10-20) siRNA using RNAimax according to the manufacturer’s instructions. Transfection was repeated 3 days later and cells were harvested for analysis at day 6.

### High-throughput Screening (HTS)

The chemical compound library was provided by the Chemical Biology Consortium Sweden (CBCS) and contained 4,126 pharmacologically active compounds from the following libraries: Prestwick, Tocris mini, Selleck tool compounds, Selleck known kinase inhibitors and ENZO tool compounds as well as 115 covalent drugs synthesized by Henriksson M. (Karolinska Institutet, Sweden). Plate and liquid handling were performed using Echo550 (Labcyte), Viaflo 384 (Integra Bioscience) and Multiflo FX Multi-Mode Dispenser (BioTek). Images were acquired by IN Cell Analyzer 2200 (GE Healthcare) with a 4x objective and quantitative image analyses were run in CellProfiler (www.cellprofiler.org) (Jones, Kang et al., 2008). Statistical analyses were carried out with Microsoft Excel and Graphpad Prism softwares. For the primary HTS screening, A549 cells were trypsinized, resuspended in culture medium containing 100ng/ml human IFNγ(Sigma-Aldrich), dispensed into 384-well plates (BD Falcon, 353962) and incubated for 24h at 37°C in a 5% CO_2_ atmosphere. The next day, compounds were added to cells achieving a final concentration of 10 μM and a DMSO volume concentration of 0.1%. Cells were incubated for 24h before staining with PE-labeled anti-human CD274 (Clone MIH1, BD Biosciences) antibody. Cells were fixed and stained with 2% formaldehyde and 2 μM Hoechst 33342, respectively. For the validation screening, MCF-7 cells were hormone-stripped by pre-culturing in SFM for 15 days with several media changes. Cells were subsequently exposed to DMSO or ethinylestradiol (EE) for 3 days, seeded in 384-well plates and incubated overnight at 37°C in a 5% CO_2_ atmosphere. The next day, the chemical library, comprising 163 compounds, including estrogens and antiestrogens, was added to the cells reaching a final concentration of 0.1, 1.0 and 10*μ*M, respectively. After 72 h of incubation, cells were stained and fixed as described above.

### Immunofluorescence microscopy

Cells were grown in 96-well imaging plates (BD Falcon, 353219) and directly stained with PE-labeled anti-human CD274 antibody (Clone MIH1, BD Biosciences) before washing with PBS and fixation with 2% formaldehyde. Subsequently, cells were permeabilized in 0.25% Triton X-100 in PBS and blocked in 3.5% BSA-PBS. Cells were stained with anti-ERα antibody (CST, 8644) diluted in blocking solution. Nuclei were stained with 2μM Hoechst 33342. Images were acquired on a Nikon eclipse Ti2 inverted wide-field microscope using a 20x 0.75 NA objective.

### Immunoblotting

Cells were lysed in RIPA buffer (Thermo Fisher Scientific) supplemented with protease and phosphatase inhibitor cocktail (Roche), sonicated for 5 min and centrifuged at 4°C, 14000 rpm for 15 min. 50μg whole-cell extracts were separated by SDS–PAGE and transferred onto Nitrocellulose membrane (Bio-Rad). After blocking in 5% milk in TBST, immunodetection was done overnight at 4°C with antibodies against PD-L1 (CST, 13684), ERα (CST, 8644) Stat1 (CST, 14994), phospho-Stat1^Tyr701^ (CST, 9167), p65 (CST, 8242), phospho-p65^Ser536^ (CST, 3033), β-actin (Abcam, ab6276) and GAPDH (Millipore Sigma, ab2302). Appropriate HRP-coupled secondary antibodies diluted in blocking solution were incubated for 1 h at room temperature. Signals were visualized by chemiluminescence (SuperSignal™ West Dura, Thermo Scientific, 34076) and acquired by an Amersham Imager 600 (GE healthcare).

### Flow cytometry

Cells were cultured as indicated and harvested using Accutase (BD Biosciences, 561527). After centrifugation, cells were stained with PE-labeled anti-human PD-L1 antibody (Clone MIH1, BD Biosciences) diluted in 2% FCS-PBS blocking solution for 45 min at 4°C. Samples were washed and immediately measured on a Guava easyCyte flow cytometer (EMD Millipore). Data were analyzed with Guava InCyte and GraphPad Prism software.

### qRT-PCR

Total RNA was isolated using the PureLink RNA mini kit (Invitrogen) according to the manufacturer’s instructions. Reverse transcription and PCR amplification were performed using TaqMan RNA-to-CT 1-Step Kit and the StepOnePlus™ real-time PCR instrument (Applied Biosystems). The following probes were used in this study: Hs01125301_m1 for *CD274*, Hs99999901_s1 for *18S*, Hs03929097_g1 for *GAPDH*, Hs01046817_m1 for *ESR1*, Hs01100353_m1 for *ESR2*, Hs00174128_m1 for *TNFA*, Hs00989291_m1 for *IFNG*.

### Cytokine analysis

MCF-7 cells were cultured in different media for indicated number of days and supernatant culture media was collected every 4 days, centrifuged and stored at −80°C until analysis. IL-6 levels of culture supernatants were determined using the LEGENDplex™ Human Inflammation Panel I (BioLegend, 740809), according to the manufacturer’s instructions. Briefly, supernatants (50μL /sample) were incubated with capture beads for 2 h at room temperature on an orbital shaker. Next, detection antibody was added and beads were incubated for 1 h at room temperature. After washing the beads, samples were measured using a BD LSRFortessa™ flow cytometer (BD Biosciences). Cytokine concentration was calculated based on a standard curve using BioLegend’s LEGENDplex™ data analysis software.

### Gene expression analysis

Next-generation RNA sequencing was performed to determine changes in gene expression between DMEM (normal), SFM (Treatment 1), fulvestrant (Treatment 2) or SFM+EE (Treatment 3) cultured MCF-7 cells. Cells were cultured during 20 days in DMEM, SFM or fulvestrant, and 20 in SFM with addition of ethinylestradiol the last 4 days. The cells were subsequently harvested and frozen at −80 °C, and the samples were processed by Eurofins Genomics Sweden AB, were the RNA was isolated and assessed for QC, and finally cDNA library preparation was performed. Illumina single read sequencing with a read length of 1x 50 bp and 30million reads per sample was performed.

### RNA sequencing data analysis

For gene set enrichment analysis (GSEA), genes in each condition were ranked based on the log2 (fold change) value between DMEM (normal) and SFM (Treatment 1) or fulvestrant (Treatment 2). Each treatment was done in triplicate. Genes enriched in each treatment were positive and genes enriched in normal conditions (DMEM) were negative. The ranked gene lists were loaded into GSEA software and tested against the gene sets of the Hallmark collection (MSigDB).

### Data availability

RNA sequencing data associated to this work are accessible at the GEO repository, under accession number GSE134938.

### T-cell mediated tumor cell killing assay

MCF7 cells were transfected with mCherry-Nucleus-7, a gift from Michael Davidson (Addgene plasmid # 55110), and a clone was selected. This MCF7 clone was then subjected to SFM, SFM with 10nM ethinylestradiol and fulvestrant treatments, as described above. T-cells were isolated 5 days prior to addition to MCF7 cells and were activated with Dynabeads^®^ Human T-Activator CD3/CD28 (Thermofisher, 11131D) and 2.5 ng/mL of recombinant human IL-2 (Thermofisher, PHC0027). T-cells were co-cultured with the mCherry-Nucleus-7 MCF7 cells in the presence of CellEvent™ Caspase-3/7 Green Detection Reagent (ThermoFisher, C10423) in 96 well plates. Images were captured every 2 hours on a MetaXpress Microscope (Molecular Devices). Total MCF7 nuclear count and caspase intensity was analysed using CellProfiler software.

### TCGA data set analysis

mRNA expression levels of *CD274* from the METABRIC breast cancer TCGA data set (Pereira, Chin et al., 2016) were retrieved from cBioportal. Only patients with tumor mRNA data were taken into consideration (n=1904). The samples were then classified as ERα+ or ERα− using the annotation included in the dataset. The expression levels present in cBioportal are automatically transformed into Z-scores for comparison purposes. Two-tailed Student’s t-test was used to assess the statistically significant differences in *CD274* mRNA expression levels between ERα^+^ or ERα^−^ patients.

### Mouse study

PyMT [FVB/N-Tg(MMTV-PyVT)634Mul/J] transgenic animals harboring breast tumors were treated with 4-hydroxytamoxifen (Sigma-Aldrich) (1.2 mg/kg/daily) in 10% ethanol in sunflower oil by oral gavage. Animals were killed in CO_2_ chamber when tumors reached the humane end point, and the tumors were fixed in 10% formalin solution and embedded in paraffin. For purification of total RNA from formalin-fixed tumor sections, RNeasy FFPE kit (Qiagen) were used following manufacturer instructions. Reverse transcription was done using SuperScriptTM IV VILOTM Master Mix (ThermoFisher Scientific) and the real time PCR was performed using Fast SYBRTM Green Master Mix (Applied Biosystems) in a 7500 Fast Real-Time PCR system (Applied Biosystems). The following primers were used for *Cd274* (FW: 5’ TGCGGACTACAAGCGAATCA and REV: 5’ GCTGGATCCACGGAAATTC) and β-actin detection (FW: 5’ GGCTCCTAGCACCATGAAGA and REV: 5’ CCACCGATCCACACAGAGTA).

### Human study and tissue

Women with a histologic diagnosis of HRPBC, for whom tissue from a distant metastasis and full medical records were available, were eligible. Patients with synchronous metastases were excluded. The study protocol was approved by the Institutional Review Board of Hospital 12 de Octubre (“Comité Ético de Investigación Clínica - Hospital 12 de Octubre”, Madrid, Spain) (Study code: 11/137), and conducted according to the principles expressed in the Declaration of Helsinki. This review board waived the need for consent since all the samples belonged to patients diagnosed of cancer before 2007. According to the Royal Act in Biomedical Research in force in Spain since 2007 (Royal Act 14/2007, July 3^rd^), the retrospective collection of archival samples belonging to patients diagnosed before 2007 do not require individual signed informed consent.

### Immunohistochemistry

For histological analyses, tissues were fixed in 10% buffered formalin (Sigma-Aldrich; St. Louis, MO, USA) and embedded in paraffin. Immunohistochemical staining with anti-PD-L1 antibody (Rabbit monoclonal (E1L3N)-Cell Signaling #13684) was performed on 2.5-μm tissue sections. Immunohistochemistry was performed using an automated protocol developed for the Autostainer Link automated slide staining system (DAKO, Agilent). All steps were performed on this staining platform using validated reagents, including deparaffinization, antigen retrieval (cell conditioning), and antibody incubation and detection. Corresponding stainings were acquired and digitalized using the AxioScan.Z1 system (Zeiss). Digitalized images were automatically analyzed with the AxioVision version 4.6.2 software (Zeiss). The percentage of PD-L1 positivity was considered as ratio of PD-L1-positive cells to total number of cells.

### Statistics

Statistical parameters and tests are reported in the Figures and corresponding Figure Legends. Statistical analysis was done using GraphPad Prism version 8.0 (GraphPad Software Inc). One-way-ANOVA was performed for all the datasets that required comparison among multiple data points within a given experimental condition.

## ACKNOWLEDGEMENTS

We would want to thank Andres J Lopez-Contreras for insightful comments on the manuscript. Research was funded by grants from the Cancerfonden foundation (CAN 2018/381) and the Swedish Research Council (VR) (538-2014-31) to OF and from the ISCIII (AES -PI16/00354; co-funded by the European Regional Development Fund) and from the Call for Coordinated Research Groups from Madrid Region, Madrid Regional Government-ERDF funds (B2017/BMD3733) to MQF.

## AUTHOR CONTRIBUTIONS

D.H. and P.M-R. contributed to most experiments and data analyses and to the preparation of the figures. C.S.H. contributed to the T-cell killing experiments. S.M. and M.Q-F helped with experiments using MMTV-PyMT animals and with human patient material. M. H. helped with the chemical screen. L.L. provided general technical support to many of the experiments. O.F-C. and J.C-P. supervised the study and wrote the MS.

