## Supplementary material for "Estrogen deprivation triggers an immunosuppressive phenotype in breast cancer cells": Figures S1-6, Table S3

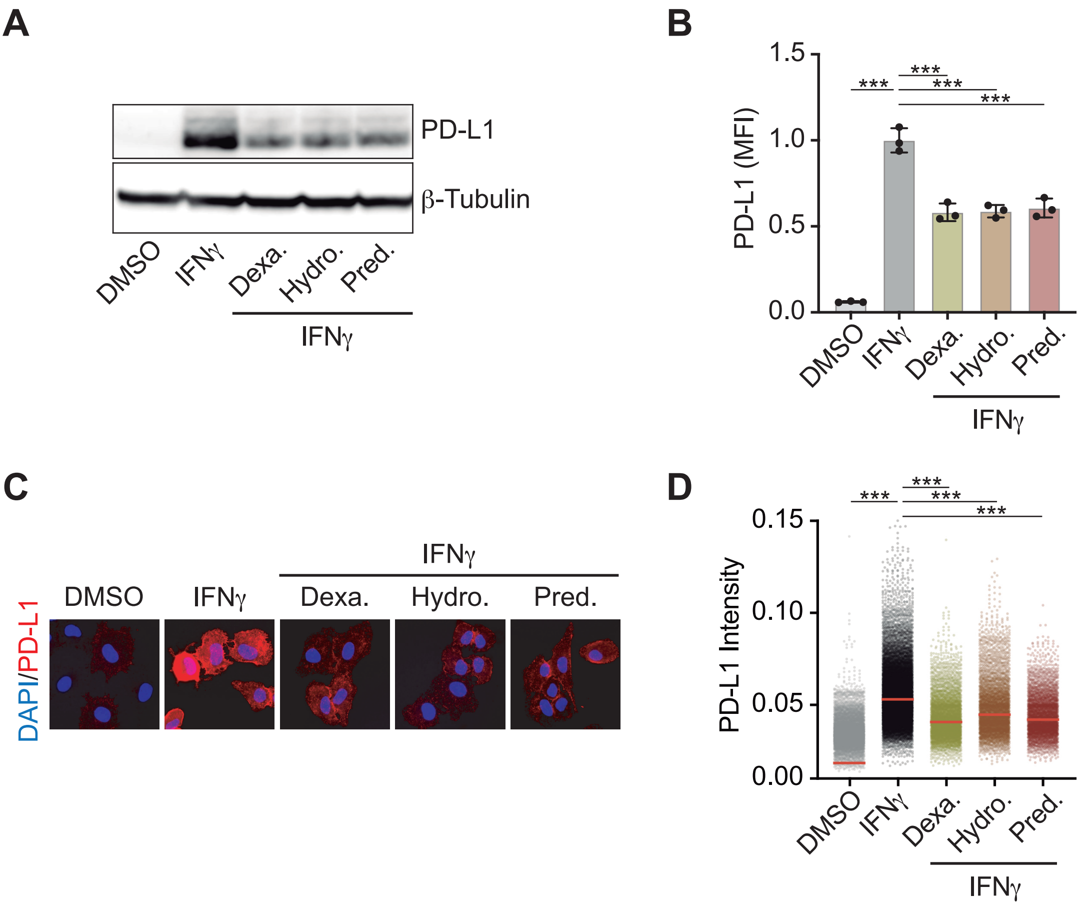
A
PD-L1
β-Tubulin
DMSO
IFNγ
Dexa.
Hydro.
Pred.
IFNγ
B
PD-L1 (MFI)
1.5
1.0
0.5
0.0
***
DMSO
IFNγ
Dexa.
Hydro.
Pred.
IFNγ
C
DAPI/PD-L1
IFNγ
DMSO
IFNγ
Dexa.
Hydro.
Pred.
D
PD-L1 Intensity
0.15
0.10
0.05
0.00
***
DMSO
IFNγ
Dexa.
Hydro.
Pred.
IFNγ

**Figure S1. Corticosteroids reduce IFNγ-induced PD-L1 expression in A549 cells A,** Western blot illustrating the levels of PD-L1 in A549 cells grown in the presence of DMSO as control or 100ng/ml IFNγ 24hrs before addition of Hydrocortisone (10μM), Prednosolone (10μM) or Dexamehtasone (10μM), for 24hrs. β-Tubulin levels are shown for loading control. **B,** Quantification of flow cytometry mediated assessment of surface PD-L1 levels in A549 after 24hrs of control or compound exposure (as in **A**). **C,** Immunofluorescence of PD-L1 (red) in A549 cells cultured in the presence or absence of IFNγ and the indicated compounds (as in **A**). **D,** After 24hrs of compound exposure (as in **A**), cells were stained with an anti-PD-L1 antibody conjugated to phycoerythrin (PE-Cy3) and fixed with formaldehyde. Nuclei were stained with Hoechst and the immunofluorescences were analyzed by high-throughput microscopy (HTM).

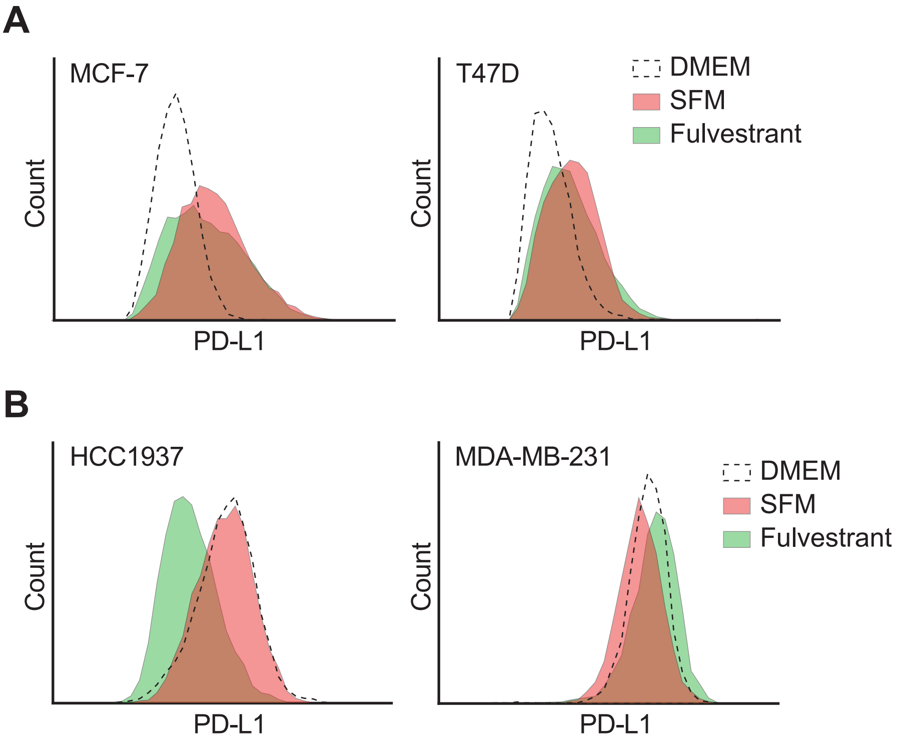
A
MCF-7
Count
PD-L1
T47D
Count
PD-L1
DMEM
SFM
Fulvestrant
B
HCC1937
Count
PD-L1
MDA-MB-231
Count
PD-L1
DMEM
SFM
Fulvestrant

**Figure S2**. **Estrogen-deprival induces PD-L1 expression in ER+ BC cells A,** Flow cytometry-mediated quantification of PD-L1 surface levels in $ER\alpha^+$ BC cell lines (MCF7 and T47D), grown in DMEM, SFM or DMEM supplemented with fulvestrant (1 μM) for 18 days. **B,** Flow cytometry-mediated quantification of PD-L1 surface levels in $ER\alpha^-$ BC cell lines (HCC1937 and MDA-MB-231), grown in normal medium (DMEM for MDA-MB-231 and RPMI 1640 for HCC1937), SFM or medium supplemented with fulvestrant (1 μM) for 18 days.

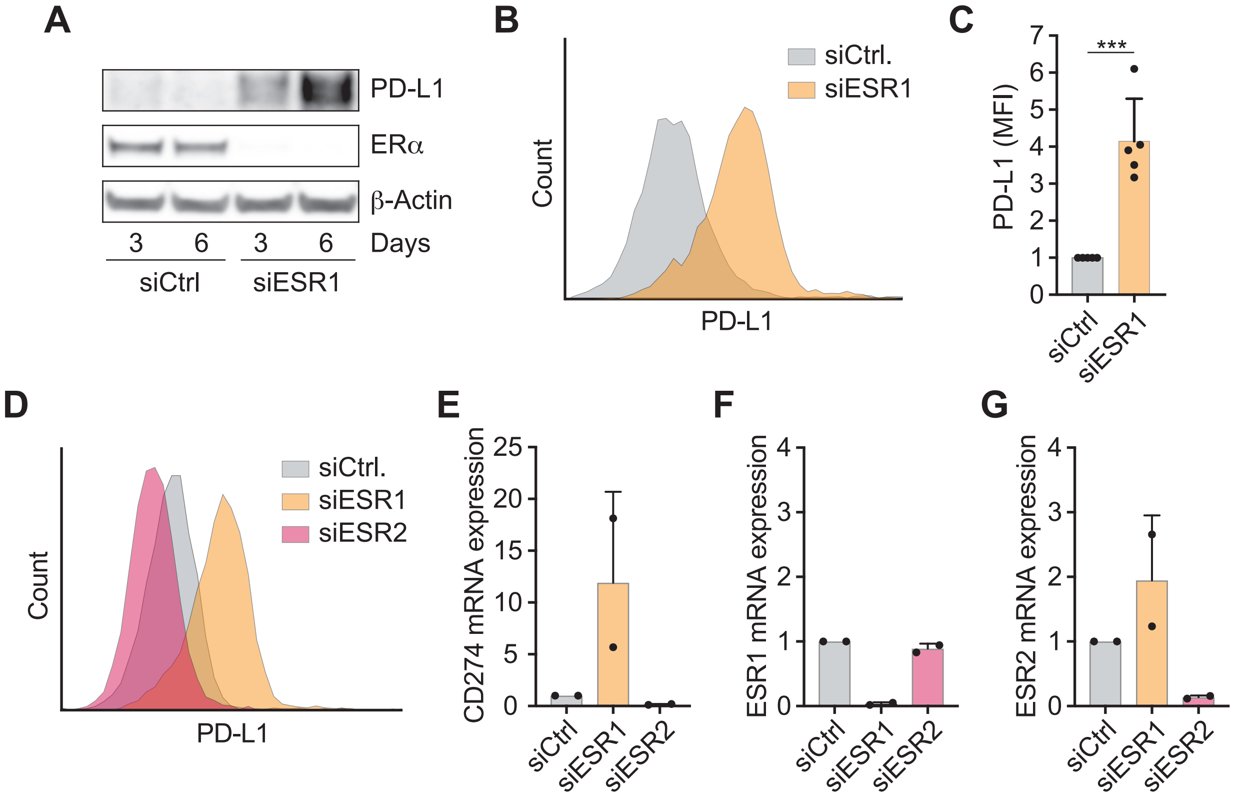
A
PD-L1
ERα
β-Actin
3 6 3 6 Days
siCtrl siESR1
B
siCtrl.
siESR1
Count
PD-L1
C
***
PD-L1 (MFI)
0 1 2 3 4 5 6 7
siCtrl
siESR1
D
siCtrl.
siESR1
siESR2
Count
PD-L1
E
CD274 mRNA expression
0 5 10 15 20 25
siCtrl siESR1 siESR2
F
ESR1 mRNA expression
0 1 2 3 4
siCtrl siESR1 siESR2
G
ESR2 mRNA expression
0 1 2 3 4
siCtrl siESR1 siESR2

**Figure S3**. **ERα, and not ERβ, suppresses PD-L1 expression in MCF7 cells** **A,** Western blot analysis of the indicated proteins in MCF7 cells treated with either 20 nM control or ERα-targeting (*ESR1*) siRNA for 3 or 6 days. β-Actin is shown as a loading control. **B,** PD-L1 membrane levels analysed by flow cytometry after treatment of MCF7 cells with the indicated siRNAs for 6 days. **C,** Quantification of 3 independent experiments as the one shown in (**B**). Data are presented as mean +SD (n=5). ***$p$<0.001. **D,** PD-L1 membrane levels as analysed by flow cytometry in MCF7 cells transfected twice with 20nM of the indicated siRNAs for 6 days. **E-G** qRT-PCR analysis of *CD274* (**E**), *ESR1* (**F**) and *ESR2* (**G**) mRNA levels in MCF7 cells transfected as in (**D**). Levels of the *18S* rRNA served as an internal control.

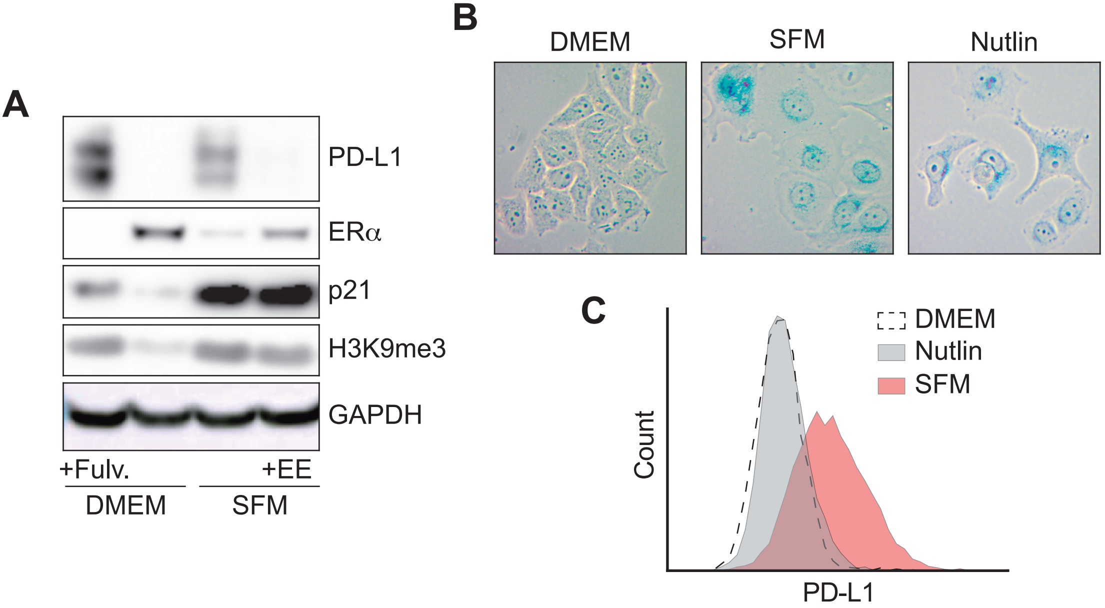
A
PD-L1
ERα
p21
H3K9me3
GAPDH
+Fulv.
+EE
DMEM
SFM
B
DMEM
SFM
Nutlin
C
DMEM
Nutlin
SFM
Count
PD-L1

**Figure S4**. **Senescence fails to upregulate PD-L1 expression in MCF7 cells** **A,** MCF7 cells were grown in DMEM, SFM or DMEM with fulvestrant (1 μM) for 17 days. Where indicated, EE (10 nM) was added for the last 3 days. Whole-cell extracts were subjected to western blot analysis of the indicated proteins. **B,** SA-β-Galactosidase assay of MCF7 cells cultured in DMEM, SFM (19 days) or Nutlin-3 (5 μM, 5 days). Representative images for each condition are shown. **C,** FACS analysis of PD-L1 membrane levels in MCF7 cells cultivated as in (**B**).

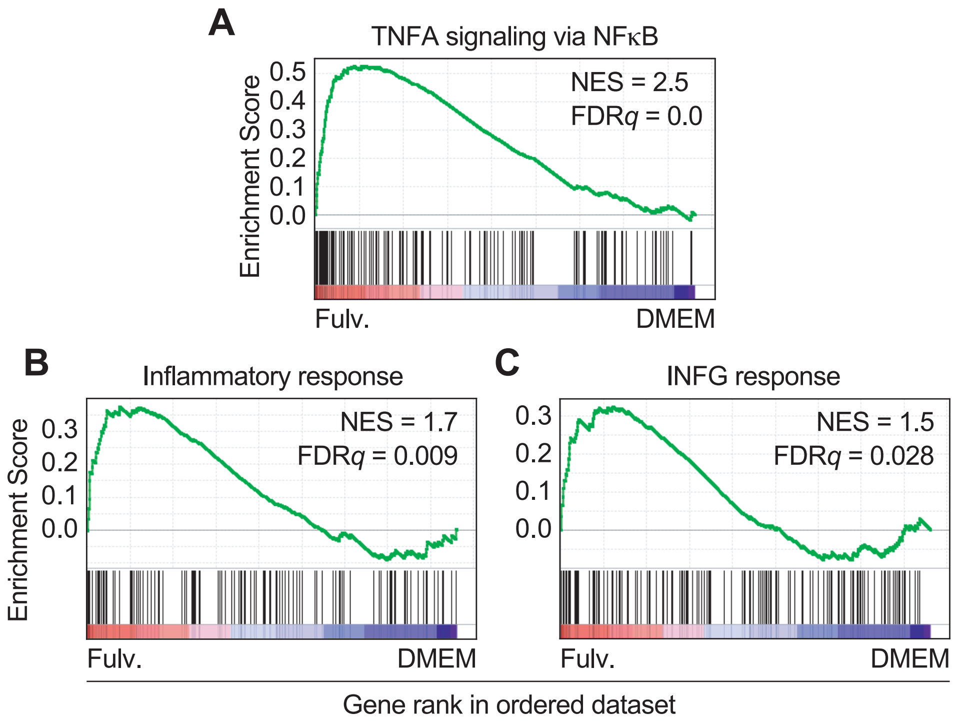
A
TNFA signaling via NFκB
Enrichment Score
0.5
0.4
0.3
0.2
0.1
0.0
NES = 2.5
FDRq = 0.0
Fulv.
DMEM
B
Inflammatory response
Enrichment Score
0.3
0.2
0.1
0.0
NES = 1.7
FDRq = 0.009
Fulv.
DMEM
C
INFG response
0.3
0.2
0.1
0.0
NES = 1.5
FDRq = 0.028
Fulv.
DMEM
Gene rank in ordered dataset

**Figure S5**. **Fulvestrant triggers an inflammatory transcriptional program in MCF7 cells A-C,** Pre-ranked GSEA on the genes from the hallmarks "TNFA signaling via NF-κB" (**A**), "Inflammatory response" (**B**) and "IFNG response" (**C**) obtained from RNAseq analysis comparing the transcriptome of MCF7 cells grown in DMEM or fulvestrant (1 μM) for 3 weeks. The heatmap representation illustrates the overall upregulation of these pathways in fulvestrant-treated MCF7 cells.

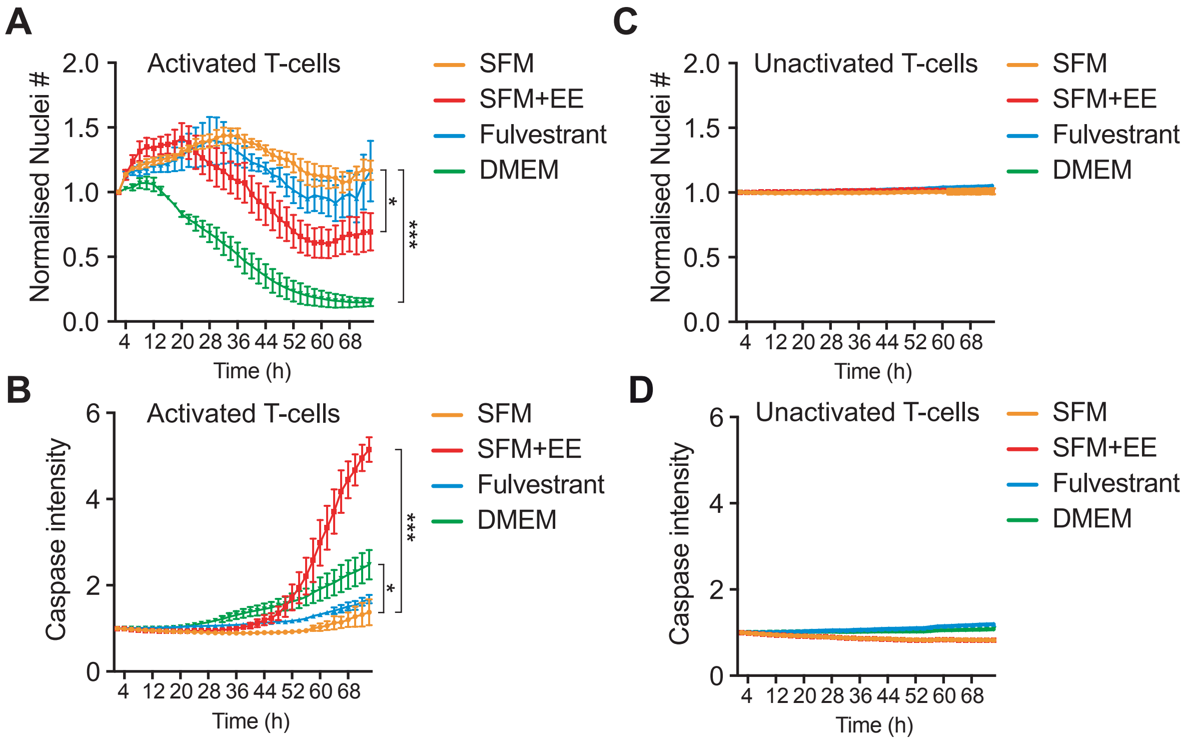
A
Activated T-cells
Normalised Nuclei #
2.0
1.5
1.0
0.5
0.0
4 12 20 28 36 44 52 60 68
Time (h)
*
***
SFM
SFM+EE
Fulvestrant
DMEM
B
Activated T-cells
Caspase intensity
6
4
2
0
4 12 20 28 36 44 52 60 68
Time (h)
***
*
SFM
SFM+EE
Fulvestrant
DMEM
C
Unactivated T-cells
Normalised Nuclei #
2.0
1.5
1.0
0.5
0.0
4 12 20 28 36 44 52 60 68
Time (h)
SFM
SFM+EE
Fulvestrant
DMEM
D
Unactivated T-cells
Caspase intensity
6
4
2
0
4 12 20 28 36 44 52 60 68
Time (h)
SFM
SFM+EE
Fulvestrant
DMEM

**Figure S6**. **Estrogen-deprival limits T-cell mediated killing of MCF7 cells A,** Live cell imaging of nuclei count from MCF7 cells after addition of activated primary T lymphocytes isolated from human peripheral blood. Nuclei counts were normalized to the value at T = 0 hrs for each condition and then each timepoint was subsequently normalized to the value of the control to which no T-cells were added. Values represent the mean ± SEM. **B,** Time-lapse microscopy of the intensity of a fluorescently labelled caspase-3/7 substrate in MCF7 cells exposed to activated primary T-cells. Values represent the mean ± SEM. **C,D** Equivalent analyses as the ones shown in (**A**) and (**B**), but where non-activated T-cells were used, illustrating that activation of the T-cells is a requisite for their cytotoxic activity against MCF7 cells. For these experiments, MCF7 cells were grown in normal media, SFM or fulvestrant (1 μM) for 2 weeks, prior to the addition of the activated T cells. Where indicated, EE (10 nM) was added for the last 3 days.

**Table S3. Clinical and demographic characteristics of the ER$^+$ BC patients used in Fig. 4.**

| **Characteristic** | **N=8** |
|---|---|
| **Age (median, range)** | 53.6 (35.1-68.2) |
| **Tumor size** | |
| T1 | 0 (0%) |
| T2 | 6 (75%) |
| T3 | 1 (12.5%) |
| T4 | 1 (12.5%) |
| **Nodal stage** | |
| N0 | 2 (25%) |
| N1 | 6 (75%) |
| **Grade** | |
| G1 | 0 (0%) |
| G2 | 7 (87.5%) |
| G3 | 1 (12.5%) |
| **Adjuvant hormonal treatment** | 0 (0%) |
| -Tamoxifen only | 3 (37.5%) |
| -Switch (Tamoxifen → Aromatase inhibitor) | 4 (50%) |
| -Aromatase inhibitor only | 1 (12.5%) |
| **Disease-free interval (years: median, range)** | 4.5 (2.84 – 15.1) |
| -Relapse during adjuvant hormonal therapy | 5 (62.5%) |
| -Relapse after completion of adjuvant hormonal therapy | 3 (37.5%) |
| **Hormonal receptors** | |
| ER and/or PR positive | 8 (100%) |
| HER2-positive | 0 (0%) |
| **Baseline Ki67** | |
| <15% | 5 (62.5%) |
| >14% | 3 (37.5%) |
